# Enhancing Therapeutic Insulin Transport from Macroencapsulated Islets Using Sub-Minute Pressure at Physiological Levels

**DOI:** 10.1101/2023.12.11.570688

**Authors:** Ella A. Thomson, Sooyeon Lee, Haixia Xu, Hannah Moeller, Joanna Sands, Rayhan A. Lal, Justin P. Annes, Ada S. Y. Poon

**Affiliations:** Department of Electrical Engineering, Stanford University, Stanford, CA 94305, USA; Division of Endocrinology, Department of Medicine, Stanford University, Stanford, CA 94305, USA; Stanford Diabetes Research Center, Stanford, CA 94305, USA; Safran ChEM-H, Stanford University, Stanford, CA 94305, USA; Division of Endocrinology, Department of Pediatrics, Stanford University, Stanford, CA 94305, USA

## Abstract

Cadaveric islet and stem cell-derived transplantations hold promise as treatments for type 1 diabetes. To tackle the issue of immunocompatibility, numerous cellular macroencapsulation techniques have been developed that utilize diffusion to transport insulin across an immunoisolating barrier. However, despite several devices progressing to human clinical trials, none have successfully managed to attain physiologic glucose control or insulin independence. Based on empirical evidence, macroencapsulation methods with multilayered, high islet surface density are incompatible with homeostatic, on-demand insulin delivery and physiologic glucose regulation, when reliant solely on diffusion. An additional driving force is essential to overcome the distance limit of diffusion. In this study, we present both theoretical proof and experimental validation that applying pressure at levels comparable to physiological diastolic blood pressure significantly enhances insulin flux across immunoisolation membranes—increasing it by nearly three orders of magnitude. This significant enhancement in transport rate allows for precise, sub-minute regulation of both bolus and basal insulin delivery. By incorporating this technique with a pump-based extravascular system, we demonstrate the ability to rapidly reduce glucose levels in diabetic rodent models, effectively replicating the timescale and therapeutic effect of subcutaneous insulin injection or infusion. This advance provides a potential path towards achieving insulin independence with islet macroencapsulation.

**One Sentence Summary:** Towards improved glucose control, applying sub-minute pressure at physiological levels enhances therapeutic insulin transport from macroencapsulated islets.

## INTRODUCTION

Type 1 diabetes (T1D) is characterized by autoimmune islet beta-cell destruction and insulin deficiency. Current treatment utilizes exogenous insulin administered *via* subcutaneous injection or infusion. Automated insulin dosing (AID) is increasingly common but still limited in efficacy and uptake^1^, particularly among racial and ethnic minorities^2^. Additionally, T1D treatment remains burdensome, costly, and associated with acute, life-threatening hypoglycemia. Individuals living with diabetes require frequent or continuous glucose monitoring, wearing a pump or administering injections, and accounting for carbohydrate ingestion and activity that alters insulin sensitivity^3^. Alternatively, islet transplantation could alleviate patient management burden and cost. In perhaps the simplest incarnation, allogeneic islets are infused into the liver *via* the portal vein^4,5^. While this approach fosters islet function and viability, providing insulin independence in the short term, transplants generally fail due to immunosuppressive toxicity and insufficiency^6^. Recently, the VX-880, which involves hepatic portal vein infusion of fully differentiated allogeneic stem cell-derived insulin-producing islet cells, found success in clinical trials, with two people achieving insulin independence^7,8^. Nevertheless, the need for immunosuppression remains a barrier to widespread deployment. To address the issue of immunocompatibility, islet encapsulation has been introduced. Nano- and micro-encapsulation effectively maximize the surface area for oxygen and nutrient transfer^9,10^. However, microcapsules struggle to achieve full immunoisolation^10–12^, and the difficulty in retrieving these transplants raises significant safety concerns.

Macroencapsulation devices provide both immune isolation and retrievability, and are among the most actively pursued transplantation approaches. Numerous immunoisolating membrane have been investigated, with select systems advancing to clinical trials, such as the βAir device^13,14^, VC-01^15^, VX-264^16^, VC-02^17,18^, and the Sernova cell pouch^19,20^. While encapsulation provides a beneficial physical barrier against host immune cell entry, this immunoprotective barrier also impedes the movement of oxygen and insulin. Relying solely on diffusion for oxygen and insulin transport has led to mostly thin and planar designs, namely islet sheets^21,22^ (Fig. 1a). For an implant with a thickness of 400 µm and a standard porous Polyethylene terephthalate (PET) membrane (10 µm thick and 10% porosity), diffusion alone would take approximately 10 minutes to transport 90% of the secreted insulin. This duration aligns with the timing of the first phase of glucose-stimulated insulin secretion (GSIS). To prevent hypoxia, islet surface densities are typically maintained at less than 500 IEQ per square centimeter^23^. Per the Edmonton protocol, the clinical therapeutic dose ranges from 5,000 to 10,000 IEQ per kg^5^. This translates to a required implant surface area of 700 to 1,400 cm^2^ for a 70-kg individual to realize efficient diffusion-driven insulin transport; an area which is not clinically practical. To make it practical, the islet surface density must be increased. The VC-02 and the Sernova cell pouch both enhance islet packing density. To prevent hypoxia, their membranes are designed with multiple large pores distributed throughout to support vascularization; however, this design necessitates the use of immunosuppression. Although increased C-peptide levels have been observed, they have not yet enabled insulin independence^17–19^.

**Fig. 1.**
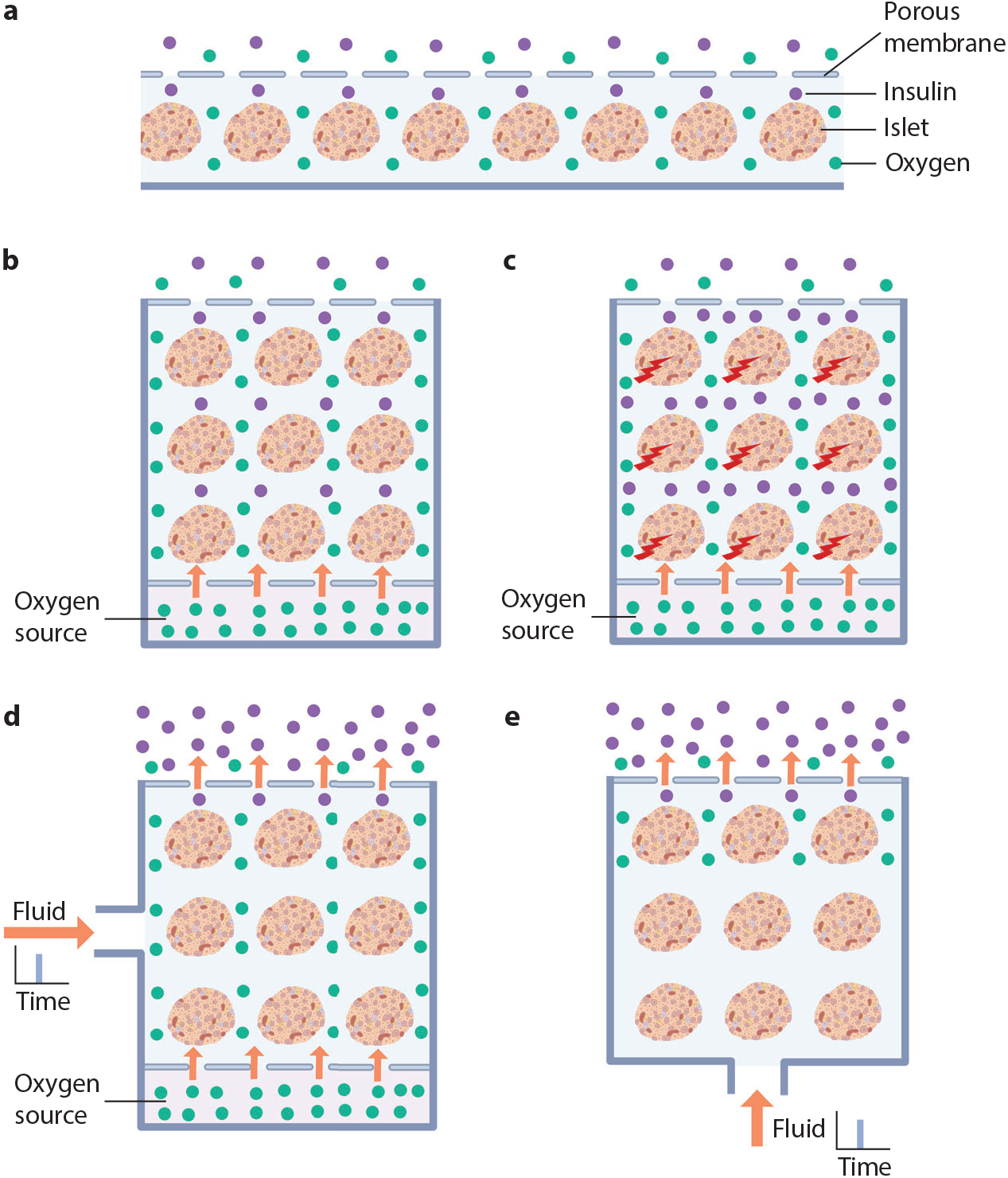
Illustration of macroencapsulation approaches including islet sheets, subcutaneous approaches with an oxygen source, and our proposed approach incorporating applied pressure. **a,** Diffusion-based methods utilize passive transport for oxygen and insulin, predominantly resulting in designs that are thin and planar due to the limitations of diffusion distances. **b,** Oxygen is either directly supplied to the macroencapsulated islets or generated *in situ*, which allows for a three-dimensional structure that increases islet surface density. However, insulin transport still primarily depends on diffusion. **c,** Cell stimulation elevates insulin levels within the cell chamber and does not overcome the diffusion limitation. **d, e,** The application of brief pressure effectively overcomes diffusion limitations in both macroencapsulation devices, with (**d**) and without (**e**) enhanced oxygenation.

To enhance islet oxygenation while maintaining immunoisolation, the physiological pressure difference between arteries and veins is utilized. This process facilitates convective transport of oxygen and essential nutrients^24–30^. High islet viability, at a density of 5,700 IEQ/cm^2^, has been demonstrated in a porcine model^30^. However, these systems are associated with substantial surgical risks. To mitigate risk and enable subcutaneous transplantation, pump-based extravascular methods are being investigated^13,14,31–34^. Among these, the βAir device, which utilizes an exogeneous oxygen supply has advanced to clinical trials but has not successfully demonstrated therapeutic glucose control or insulin independence^13,14^. Additionally, other techniques supporting subcutaneous transplantation with enhanced oxygenation include prevascularization of graft sites^35^ and *in situ* oxygen generation methods such as hydration of solid peroxide^36,37^, recycling of carbon dioxide^38^, electrolysis^39,40^, and microalga-based photosynthesis^41^. In these subcutaneous approaches, insulin transport predominantly relies on diffusion as illustrated in Fig. 1b. For instance, with islets encapsulated within a PET membrane spanning 25 cm^2^, the islet surface density must be increased by at least 28 times per the Edmonton protocol. If the islet volume density remains constant, the thickness of the cell chamber must also increase 28-fold. Consequently, the diffusion time for insulin would increase proportionally, meaning diffusion alone would require approximately ∼5 hours to transport 90% of the insulin secreted within the cell chamber. Furthermore, cell stimulation only leads to an increase in insulin levels within the cell chamber^42,43^ and is not able to overcome the diffusion limitation, as illustrated in Fig. 1c.

Thus far, macroencapsulation strategies have primarily focused on enhancing cell viability to achieve high islet surface density. However, the discussion often overlooks the diffusion limitation for insulin transport across the immunoisolating membrane. This gap poses significant challenges in achieving effective on-demand insulin delivery and maintaining physiological glucose regulation—both essential for homeostasis. Regrettably, this crucial limitation has been overshadowed by extensive research that has “validated” islet encapsulation methods in mice, demonstrating improved or normalized glucose tolerance but without a corresponding increase in insulin and/or C-peptide release. These studies often neglect the well-documented effect of glucose effectiveness—the ability of glucose to manage its own disposal and hepatic glucose output^34,35^. This effect is more pronounced in rodents than in humans^44–46^, with the magnitude of this difference being at least twice as significant^47^. In streptozotocin (STZ)-induced diabetic rodents, simply implanting an insulin pellet, that supplies only basal insulin, along with glucose effectiveness, suffices to normalize glucose tolerance tests^44^. Similarly, the βAir device, when implanted in diabetic rodents, resulted in normal glucose tolerance test outcomes, yet data reflecting the release of insulin or C-peptide are not provided^31^. Achieving normal glucose tolerance test results, particularly in rodents, does not affirm an adequate insulin transport needed for human efficacy.

To harness the clinical potential of macroencapsulated cell-based therapy, an additional driving force is required to enhance and regulate insulin transport across the immunoisolating membrane, as illustrated in Fig. 1d, e. To enable subcutaneous transplantation, we propose an extravascular pump-based method to surpass the barriers of diffusion. Our mathematical and experimental results show that applying pressure for a sub-minute duration at a level akin to physiological diastolic blood pressure is sufficient to rapidly and effectively transport insulin through encapsulating membranes. This method replicates the delivery dynamics of native islets or external insulin infusions and facilitates repeated bolus insulin delivery. By harnessing a second driving force, we acutely restored euglycemia in mice with complete or near-complete lack of endogenous insulin production within a timeframe unattainable through diffusion alone.

## RESULTS

### Order-of-magnitude analysis of pressure duration and magnitude on insulin transport across porous membranes

We consider the simultaneous effect of pressure (*p*) and concentration (*c_s_*) gradients on the transport of insulin across a porous membrane. The insulin (solute) flux, *J_s_*, is the combined effect of the solute diffusional flow and the driving force from the volumetric solvent flux, *J_v_*. The solute diffusional flow −*D_s_*∇*c_s_* is driven by the insulin concentration gradient where *D_s_* is the insulin diffusion coefficient. The volumetric solvent flux is a pressure-driven flow; this flow in turn carries the insulin to flow through the membrane resulting in an insulin flux of *J_v_c_s_* = −*L_p_h*∇*p* ⋅ *c_s_* where *L_p_* is the filtration coefficient and ℎ is the membrane thickness. Here, we are interested in finding a simple expression for the ratio of the insulin flux with and without the applied pressure in steady state to comprehend the multiplicative effect of applied pressure relative to diffusion alone for insulin transport. This expression is given by

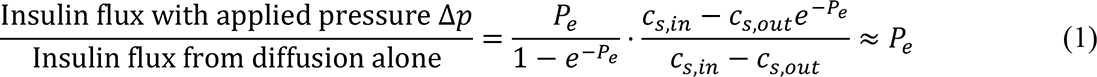

where *c_s,in_* and *c_s,out_* are the insulin concentration inside and outside the encapsulation respectively. In the ratio, *P_e_* is the axial Peclet number defined by 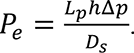 When *P_e_* is large, the ratio in Eq. 1 approximates to *P_e_*; that is, the Peclet number alone reveals the multiplicative effect of applied pressure relative to diffusion for insulin transport.

For insulin transport where membrane pore radius (*rp*) is much greater than molecular radius of insulin (*rs*), we apply the Hagen-Poiseuille Law and Strokes-Einstein Equation to obtain

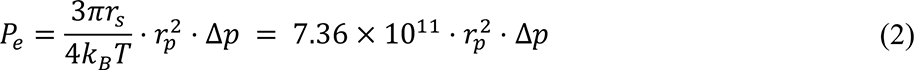

where *k_B_* is the Boltzmann constant and *T* is the absolute temperature. Here, we showed that the multiplicative effect of applied pressure compared to diffusion alone on insulin transport is primarily determined by two critical factors: the membrane’s pore radius and the magnitude of the applied pressure. For typical insulin transport membranes with pore radii around the sub-micrometer scale, this ratio is approximately 0.1Δ*p*. Thus, applying a pressure on the order of 10 kPa (comparable to normal human diastolic blood pressure, which is around 10.7 kPa) leads to nearly three-orders-of-magnitude increase in steady-state insulin flux. This is calculated as 0.1Δ*p* = 0.1 × 10^4^ = 10^5^. During transient transport, this multiplicative increase in insulin flux translates into a corresponding reduction in transport time. Specifically, a transport process that would take 5 hours *via* diffusion alone can be reduced to less than a minute.

### Validation of order-of-magnitude analysis on insulin transport with and without brief pressure

We performed numerical simulations of insulin transport across a membrane area in the range of square centimeters. Our findings revealed that relying solely on diffusion required over 2 hours to transport over 90% of encapsulated insulin (Fig. 2a). However, with the application of an 11-kPa pressure, transport time was markedly reduced to approximately 10 seconds (Fig. 2b), thereby affirming conclusions from our earlier theoretical analysis. Although our theoretical analysis used 10 kPa as an order-of-magnitude reference, we opted for 11 kPa in our simulations and experiments to more closely approximate the normal human diastolic blood pressure of about 10.7 kPa. This pressure was generated by our custom-designed piezoelectric micropump system, which effectively mimics normal human diastolic conditions.

**Fig. 2.**
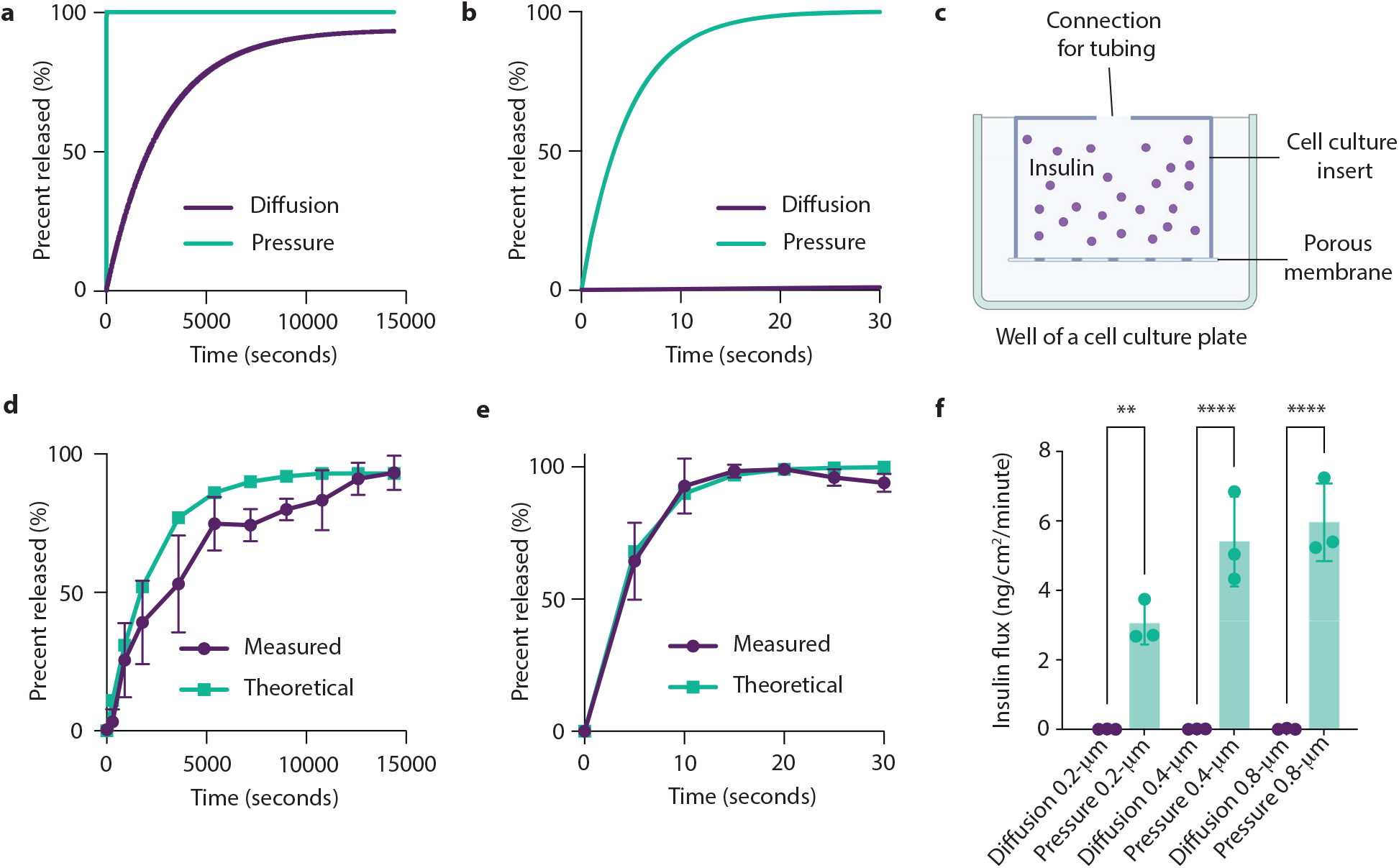
Validation of the effect of brief pressure application on insulin transport across a porous membrane through numerical simulations and experimental studies. **a, b,** Numerical simulations reveal the prolonged transport time required for diffusion-only transport (**a**), contrasting with significantly reduced transport time with the introduction of an 11-kPa pressure (**b**). Notably, (**b**) provides a zoomed-in view of (**a**) during the initial 30 seconds, emphasizing the rapid response facilitated by pressure application. **c,** Experimental setup for measuring insulin levels within and outside the porous membrane-contained space. **d**, **e**, Empirical measurements of insulin transport under diffusion-only condition (**d**) and pressure-driven transport (**e**) as compared with theoretical calculations for a membrane with 0.4 μm pore diameter. **g**, Empirical measurement of insulin flux across membranes with pore diameters of 0.2 μm (9.4% porosity), 0.4 μm (12.6% porosity), and 0.8 μm (15.1% porosity), shedding light on the relationship between pore size and insulin transport efficiency. \*\**P* < 0.01, \*\*\*\**P* < 0.0001.

Subsequently, we conducted experimental studies using a porous PET membrane with a 0.4-µm pore diameter and 12% porosity, commonly used for encapsulation^48–50,42,43^. These experiments were conducted in a 12-well format, using encapsulated cell culture inserts (Fig. 2c and Supplementary Fig. 1). Pressure was applied using a piezoelectric micropump, maintaining an average pressure of ∼11 kPa at the membrane (Supplementary Fig. 2) with no observed backflow. The empirical findings corroborated the simulation results (Fig. 2d, e), consistent across a range of pore sizes (Fig. 2f). For pore sizes ranging from 0.2 to 0.8 µm (equivalent to *rp* in Eq 2 in the range of 0.1 to 0.4 µm), the introduction of brief pressure enhanced flux by an average of approximately 400 times compared to diffusion alone. Therefore, a brief application of pressure akin to normal human diastolic blood pressure markedly shortened the time required for transporting encapsulated insulin, from several hours *via* diffusion alone to less than a minute.

### *In vitro* validation of repeated insulin bolus delivery from encapsulated R7T1 pseudoislets and human islets

Building upon the aforementioned findings, we hypothesized that (1) the encapsulation membrane hinders diffusion-dependent insulin delivery from secretagogue-stimulated β-cells; and (2) the application of a brief pressure can generate insulin boluses independently of cell stimulation, such as glucose-stimulated insulin secretion (GSIS) and KCl-triggered insulin secretion. To investigate these hypotheses, we used the setup illustrated in Fig. 1e. We utilized the R7T1 β-cell pseudoislets (Fig. 3a) and characterized them by measuring insulin secretion under both proliferating and growth arrested conditions, as well as assessing the expression of ZnT8, insulin, and PDX1 (Supplementary Fig. 3). Furthermore, we examined how the number of R7T1 pseudoislets affects insulin secretion, validating an approximately linear correlation between pseudoislet count and insulin release (Supplementary Fig. 4). In our experiments, approximately 800 pseudoislets (∼685 IEQ) were encapsulated within porous PET (200 µL, 2284 IEQ/cm^2^) using the experimental setup shown in Fig. 3b.

**Fig. 3.**
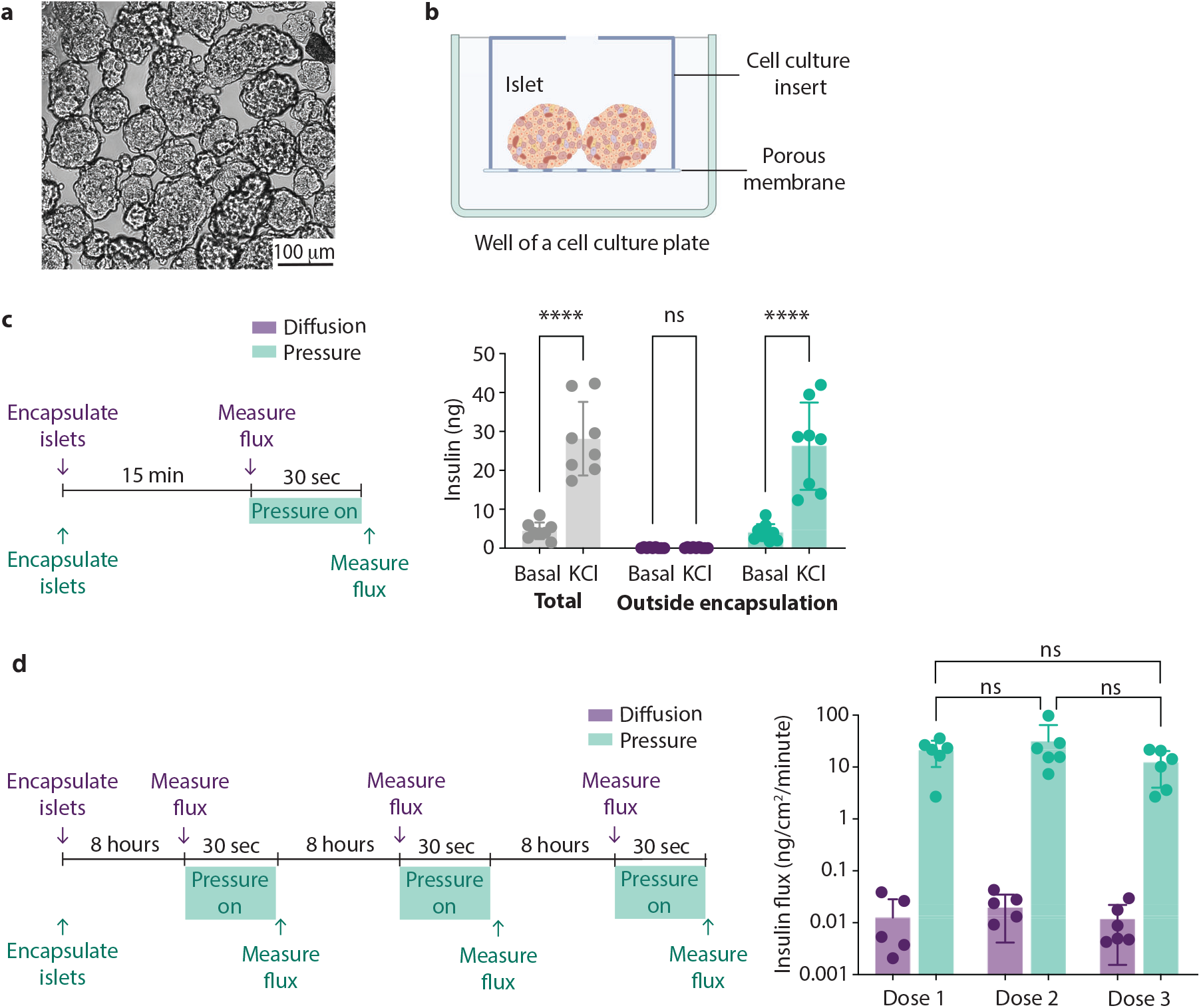
*In vitro* validation of repeated insulin bolus delivery from macroencapsulated R7T1 pseudoislets. **a**, Brightfield image of R7T1 pseudoislets. **b**, Experimental setup for measuring insulin levels released from encapsulated R7T1 pseudoislets within and outside the porous membrane-contained space. **c**, Experimental procedure for assessing KCl-stimulated and basal insulin secretion from encapsulated R7T1 pseudoislets with and without applied pressure. Measurements include total insulin released (both inside and outside of encapsulation), and insulin transported to the outside of encapsulation by diffusion and applied pressure for KCl-stimulated and basal R7T1 pseudoislets. **d**, Comparison of insulin flux between diffusion and pressure driven delivery for basal R7T1 pseudoislets over three consecutive 8-hours intervals. \*\**P* < 0.01, \*\*\*\**P* < 0.0001.

To test the first hypothesis, encapsulated pseudoislets were exposed to either basal [5 mM glucose] or stimulatory [40 mM KCl] culture conditions. As anticipated, a 15-minute KCl treatment significantly increased total secreted insulin (*p* < 0.0001; Fig. 3c). We selected the 15-minute time point to approximate the recommended pre-meal insulin bolus injection time in individuals with T1D. When solely relying on diffusion, KCl treatment failed to significantly elevate insulin levels outside of encapsulation in comparison to basal levels (*p* > 0.9; Fig. 3c). In contrast, the application of a 30-second 11-kPa pressure led to a substantial increase in insulin levels outside the encapsulation (*p* < 0.0001; Fig. 3c). These findings clearly demonstrated that KCl exposure effectively triggered insulin secretion from pseudoislets. However, achieving a corresponding elevation of insulin outside the encapsulation, on a physiologically relevant time scale, necessitated the application of a secondary driving force. These results align with previously reported observations of limited insulin release on a physiological time scale from encapsulated, electrically stimulated cells^42^.

To test our second hypothesis, we compared insulin efflux driven solely by diffusion against that facilitated by brief pressure applied to encapsulated pseudoislets after 8 hours of islet encapsulation. A 30-second 11-kPa pressure application resulted in nearly 3 orders of magnitude higher insulin flux compared to diffusion alone (Fig. 3d). This unequivocally demonstrated the ability of brief pressure to enable rapid bolus insulin delivery within a sub-minute timeframe, irrespective of cell stimulation.

In assessing potential adverse effects of this brief pressure on β-cell health and basal secretion function, we administrated two additional pressure-driven doses at 8-hour intervals. Encouragingly, no decrement in insulin release was observed across consecutive doses (Fig. 3d). Moreover, our investigation into the viability of encapsulated primary human islets *in vitro* over a 5-day period revealed comparably preserved viability with both diffusion and pressure-based insulin delivery methods (Supplementary Fig. 5). The applied pressure level aligns with physiological norms and is maintained for only a very short duration, resulting in a slow flux and ensuring safety.

Taken together, these findings indicate that a brief application of pressure facilitates repeated and consistent bolus insulin delivery independent of beta-cell stimulation. Importantly, this method does not negatively impact the function of R7T1 pseudoislets or primary human islets. Such a property holds significant advantages for the advancement of cell-based therapies, as it circumvents previously (qualitatively) described insulin transport delays^51^.

### *In vivo* validation of insulin bolus delivery using wild type mice

After confirming our findings through *in vitro* validation, we proceeded to conduct a comparative *in vivo* analysis. Subcutaneous implantation of encapsulated mouse islets was performed in wild type mice (C57BL/6J) (Fig. 4a). The implants used in all *in vivo* experiments elicited an anticipated foreign body response (Supplementary Fig. 6) at two weeks, consistent with previously described implantable devices^52^.

**Fig. 4.**
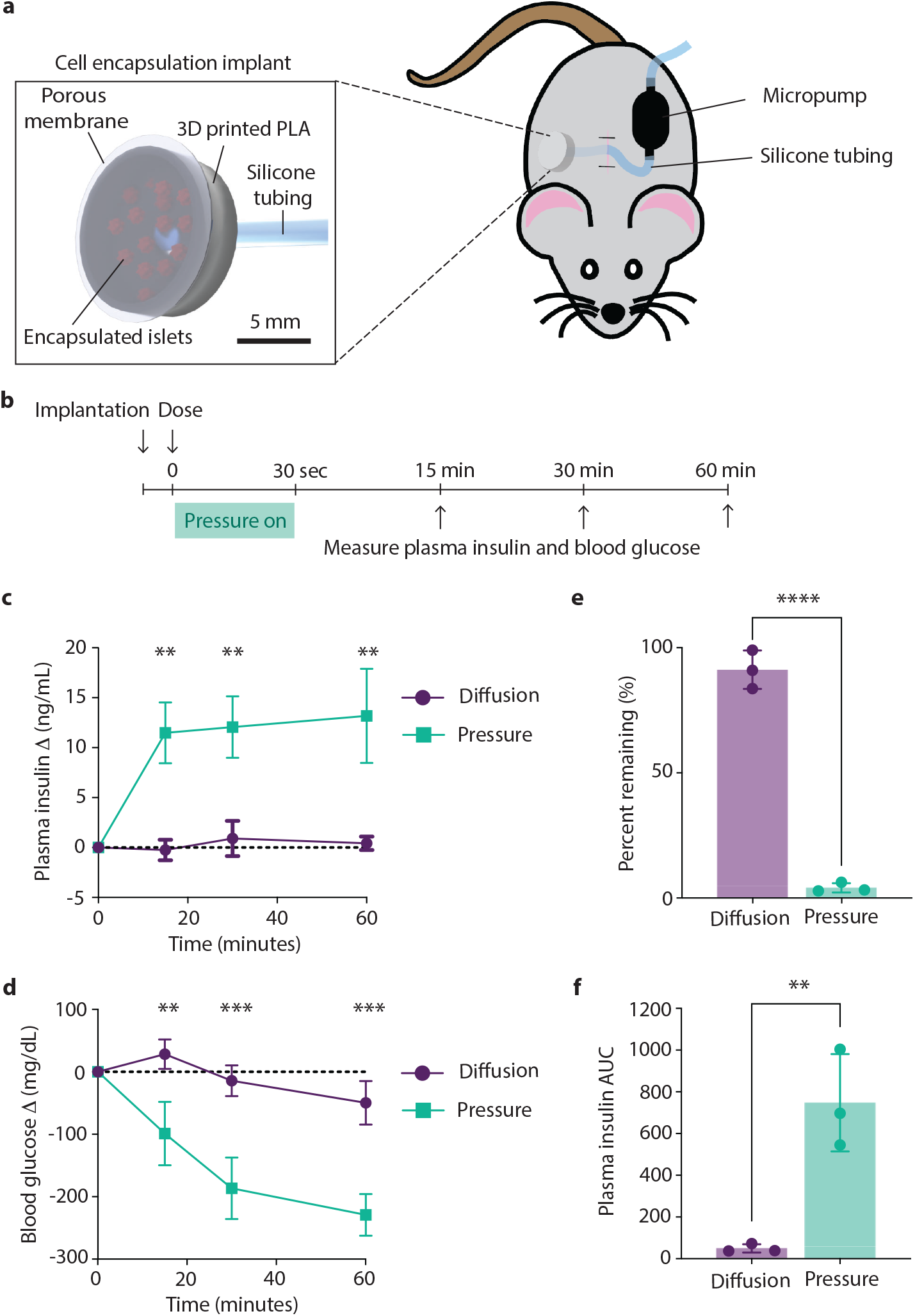
Comparative analysis of pressure-driven *versus* diffusion-based delivery from macroencapsulated mouse islets in WT mice. **a**, An illustration depicting an anesthetized mouse with a cell encapsulation implant and a micropump connected *via* silicone tubing. PLA, polylactic acid. **b**, Experimental timeline: insulin was administered almost immediately after implantation and a ∼11-kPa pressure was applied for 30 seconds for pressured-treated mice. Blood glucose levels were measured, and blood samples were collected at 0, 15, 30, and 60 minutes after initial insulin dosing. **c**, Change in blood insulin levels observed in mice treated with either diffusion or applied pressure, *n* = 3/group. **d**, Change in blood glucose levels observed in mice treated with either diffusion or applied pressure, *n* = 3/group. **e**, Percentage of dosed insulin remaining in the implant after 90 minutes, *n* = 3/group. **f**, Blood insulin AUC in mice treated with either diffusion or applied pressure, *n* = 3/group. \*\**P* < 0.01, \*\*\**P* < 0.001, \*\*\*\**P* < 0.0001.

Pressure-based dosing was performed immediately after device implantation. Following anesthesia, the average starting blood glucose was 286 mg/dL. To ensure accurate assessment of glucose effectiveness, we monitored both blood insulin and blood glucose levels post-implantation and dosing (Fig. 4b). A 30-second application of approximately 11-kPa pressure was employed for insulin bolus delivery from a 200-µL implant. Blood insulin levels in mice subjected to pressure-driven dosing were 35-fold higher than those treated with diffusion after 30 minutes (Fig. 4c). Consequently, blood glucose levels decreased by an average of 229 mg/dL with pressure-driven dosing compared to 47 mg/dL with diffusion at 60 minutes (Fig 4d). Analysis of the encapsulation device revealed that approximately 90% of the insulin secreted by the mouse islets remained within the device following 90 minutes of diffusion-based delivery, whereas less than 10% remained after pressure-driven delivery (Fig. 4e). Moreover, the blood insulin under the curve (AUC) was approximately 32-fold higher in animals receiving pressure-driven *versus* diffusion-based delivery (Fig. 4f). These results demonstrate the necessity of pressure to generate a physiologically relevant insulin bolus *in vivo*.

### Restoring euglycemia in diabetic mice using subcutaneous implant and pressure-driven insulin dosing

To assess the translational potential of pressure-driven insulin dosing, we conducted short-term *in vivo* transplantation experiments using encapsulated mouse and human islets in STZ-induced diabetic mice. Dosing was initiated immediately after device implantation. Mice dosed with applied pressure exhibited a significant increase in blood insulin levels, averaging 5.4 ng/mL for mouse islets and 7.6 ng/mL for human islets (xenogeneic transplantation) within 30 minutes. In contrast, animals treated with diffusion alone experienced a much lower increase in blood insulin levels during the same timeframe, averaging 0.04 ng/mL for mouse islets and 0.38 ng/mL for human islets. By 60 minutes, blood glucose levels decreased by an average of 368 mg/dL for mouse islets and 426 mg/dL for human islets with pressure-driven dosing, compared to an average decrease of 30 mg/dL and 57 mg/dL, respectively, with diffusion (Fig. 5a, b right). Notably, mice treated with pressure-driven dosing achieved euglycemia within 60 minutes, a result not achieved through diffusion-dependent insulin delivery from encapsulated islets. While diffusion alone did not achieve bolus delivery, a small impact on blood insulin was observed by 60 minutes (*p* = 0.03 for human islets; Fig. 5b). These findings collectively demonstrated that applying brief pressure overcame limitations posed by diffusion, enabling efficient bolus insulin delivery.

**Fig. 5.**
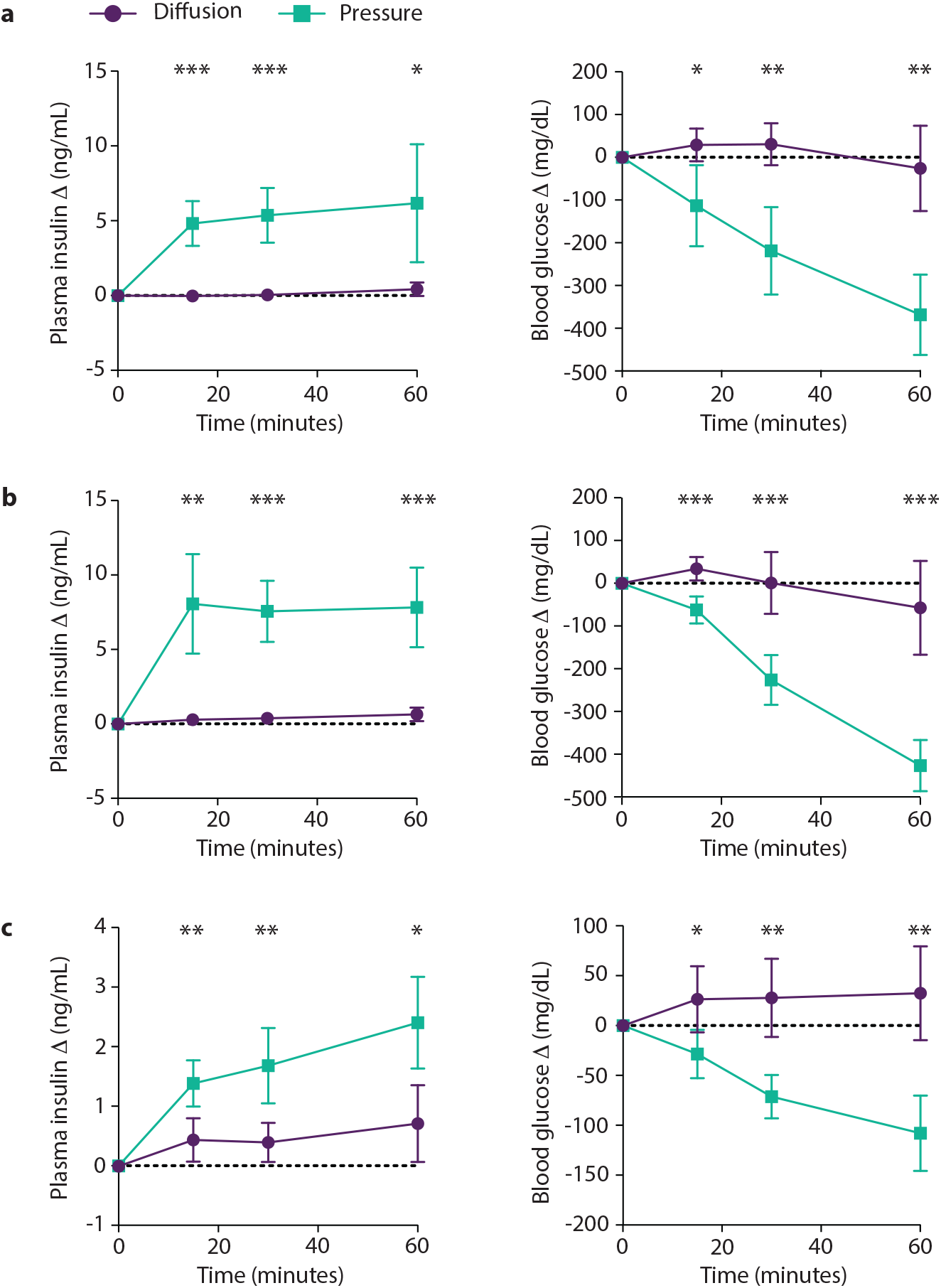
Restoring euglycemia in diabetic mice via pressure-driven insulin dosing. **a** to **c**, Changes in plasma insulin and blood glucose levels in mice subjected to either diffusion-based (purple) or pressure-driven (green) insulin release from encapsulated islets. Specific islet types used are (**a**) mouse islets, (**b**) human islets, and (**c**) R7T1 pseudoislets. *n* = 5/group. \**P* < 0.05, \*\**P* < 0.01, \*\*\**P* < 0.001.

Building on these promising results, we subsequently explored the potential of a renewable and scalable insulin source, namely reversibly transformed mouse R7T1 pseudoislets that can be proliferated as required, for managing blood glucose levels in STZ-induced hyperglycemic mice. Using pressure-driven dosing, we observed a significant reduction in blood glucose levels, with an average decrease of 108 mg/dL at 60 minutes. Conversely, when employing diffusion-based treatment, blood glucose levels actually increased by an average of 32 mg/dL, possibly due to glucose diffusion from the encapsulated cell media (Fig. 5c). This finding aligns with the modest glucose rise depicted in Fig. 4d. Our findings suggest that pressure-based dosing enhanced glucose control, offering the potential utility of a readily available biologic (R7T1 beta-cells) to traditional sources such as harvested mouse islets, human islets, or stem-cell-derived pancreatic cells.

### Enhanced glycemic control in diabetic mice using repeated pressure-driven insulin delivery with variable durations and intervals

To ascertain the clinical viability of pressure-based insulin dosing, we examined its immediate impact on blood glucose regulation in diabetic mice via repeated bolus administrations (Fig. 6a). Approximately 400 mouse islets (∼480 IEQ, 1600 IEQ/cm^2^) were encapsulated in a 200-µL implant. Within 30 minutes of the initial pressure-driven bolus, there was a notable decrease in blood glucose levels, averaging 310 mg/dL (Fig. 6b and Supplementary Fig. 7). In contrast, mice receiving insulin through passive diffusion experienced an average increase of 41 mg/dL in blood glucose levels. Sham controls showed a modest decrease of 21 mg/dL, confirming that the significant drop in glucose observed in the experimental group was attributable to the administered insulin. The kinetics of this response closely matched the pharmacokinetic profile of subcutaneously injected regular insulin^53^. Subsequent administration of a second dose 6 hours later reaffirmed these results, with pressure delivery achieving a consistent glucose reduction of approximately 270 mg/dL, while diffusion resulted in a slight increase of 32 mg/dL (Fig. 6c). The total AUC for delta blood glucose significantly favored the pressure-treated group, showing a reduction to −180,560 ± 35,003, compared to −18,549 ± 11,338 for the diffusion-treated mice (Fig. 6d). Notably, the animals were free to behave naturally between doses, highlighting the method’s robustness and reliability. By the end of the experiment, 6 hours post-final dose, all animals had returned to hyperglycemia (521 ± 35 mg/dL), underscoring that the transient euglycemia was induced by the pressure-delivered doses and not by endogenous insulin production.

**Fig. 6.**
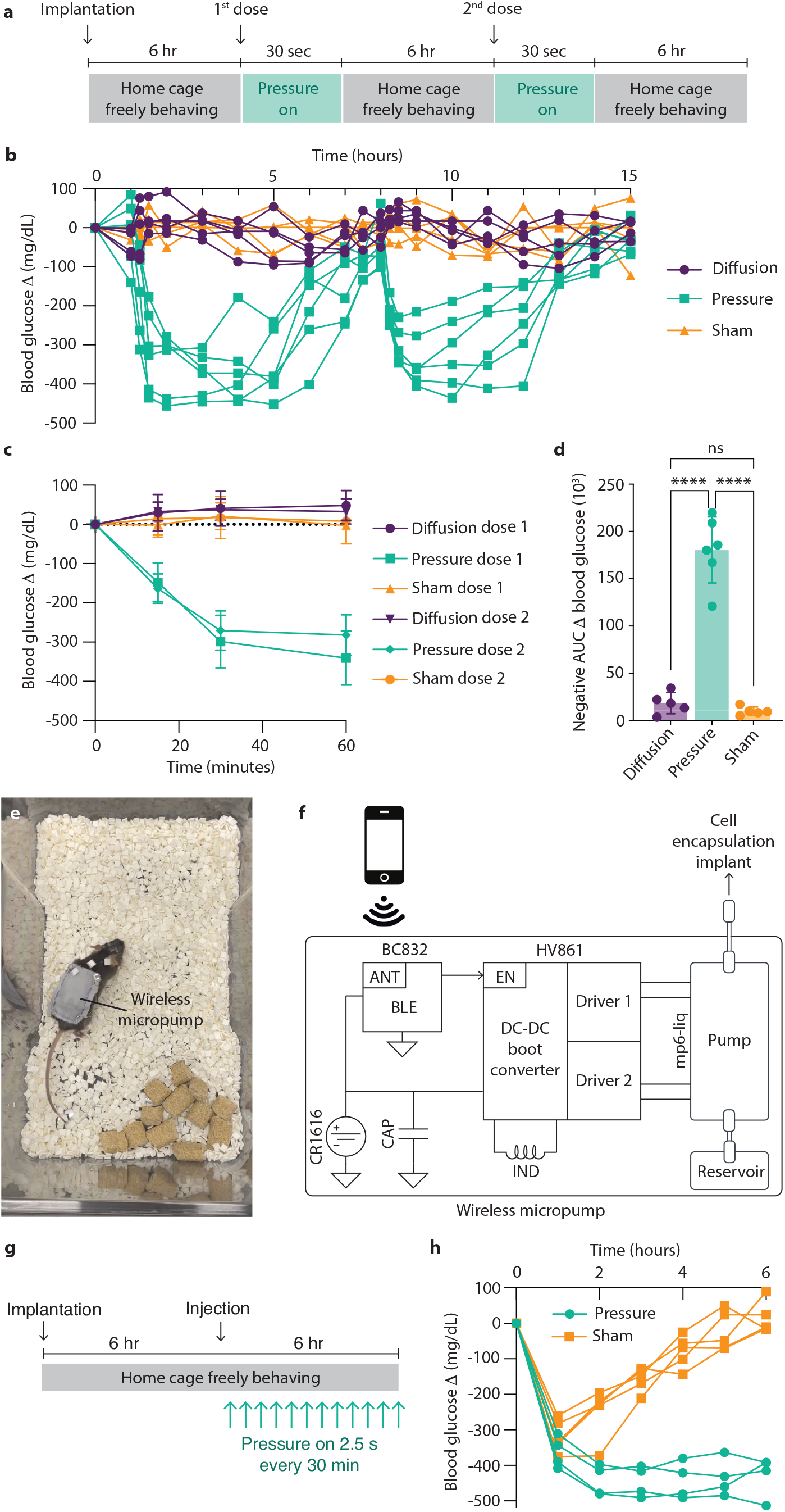
Comparing glycemic control in diabetic mice with repeated pressure-driven insulin delivery at varied durations and intervals. **a**, Schedule for insulin boluses experiments. **b**, Changes in blood glucose levels in mice treated via diffusion, applied pressure, or as sham transplanted controls; *n* = 5 per condition for diffusion and sham, *n* = 6 for pressure. **c**, Changes in blood glucose levels within 60 minutes after administering the first and second doses; *n* = 5 per condition for diffusion and sham, *n* = 6 for pressure. **d**, AUC of change in blood glucose levels for treatments by diffusion, applied pressure, and sham; *n* = 5 per condition for diffusion and sham, *n* = 6 for pressure. **e**, Photograph of the wireless micropump system on a mouse. **f**, Schematic of the wireless micropump system, featuring a high-voltage dual driver (part no. HV861), a Bluetooth transceiver module with an integrated antenna (ANT), a piezoelectric micropump (part no. mp6-liq), and a coin battery (CR1616). **g**, Schedule for basal dosing experiments. **h**, Changes in blood glucose levels in mice treated with Humalog injection or Humalog injection combined with pressure-based dosing; n = 4 for pressure, n = 5 for sham. \*\*\*\**P* < 0.0001.

Next, we demonstrated the versatility of pressure-based insulin delivery *via* adjustable dosing duration and intervals through the development of a wireless, wearable micropump system. This system facilitates repeated pressure-based dosing in freely behaving mice, thereby eliminating the need for tethering or repeated anesthesia (Fig. 6e and Supplementary Movie 1). It comprises a piezoelectric micropump, a dual-channel pump driver, a Bluetooth (BLE) controller, and a 1-mL reservoir, all powered by a single coin-cell battery (Fig. 6f and Supplementary Fig. 8). To demonstrate utility, we reconfigured the 30-second dosing schedule from every 6 hour to even shorter 2.5-second intervals every 30 minutes, while keeping the total pressure duration unchanged (Fig. 6g). An initial subcutaneous insulin injection of 0.05 units was administered to lower blood glucose levels, followed by pressure-driven dosing. The total AUC for blood glucose with pressure-based dosing reached –2556, a significant improvement over the –804 recorded for controls (Fig. 6h and Supplementary Fig. 9; *p* < 0.001). Compared to the above bolus delivery, smaller more frequent pressure-based dosing also resulted in a substantial reduction in blood glucose AUC, demonstrating short-term utility for maintaining euglycemia in mice with severe diabetes.

## DISCUSSION

For several decades, immunoisolated macroencapsulated islet transplantation has held significant unrealized promise to transform the treatment of T1D. To date, macroencapsulation strategies have generally focused on enhancing cell viability to maximize islet surface density. However, these approaches have not adequately addressed the critical challenges associated with insulin transport across the immunoisolating barrier. Often, the kinetics of insulin transport may have been under-apreciated^51,54^. It is crucial to recognize that devices relying solely on diffusion for insulin delivery may not consistently achieve effective nutrient-responsive insulin dosing.

To fully realize the clinical potential of macroencapsulated cell-based therapy, an additional mechanism to enhance insulin transport is necessary. While physiological blood pressure has been effectively utilized to enhance oxygen transport, this study explores whether a brief application of similar pressure levels could significantly improve insulin transport. Specifically, we aim to mimic the timescale and therapeutic effect of subcutaneous insulin injections or infusions. We analyzed one-dimensional flow across a porous membrane and derived an expression to compare the enhancement of insulin transport by applied pressure over diffusion alone. Critically, the enhancement depends linearly on both the pore area and the level of applied pressure. With typical pore sizes used in immunoisolation and pressures equivalent to normal human diastolic blood pressure (∼10.7 kPa), the improvement in transport rates is nearly 1000-fold. This promising theoretical insight suggests that transport times could be significantly reduced, potentially overcoming the aforementioned limitations of diffusion in insulin transport.

To validate our theoretical insight, we performed simulation to verify the substantial order-of-magnitude enhancement, followed by *in vitro* experiments using R7T1 pseudoislets. These experiments demonstrated that a brief, 30-second application of physiological-level pressure (∼11 kPa) was sufficient to reliably produce repeated insulin bolus delivery. To assess the therapeutic effect, we developed an implant based on the simplified setup illustrated in Fig. 1e, where pressure is directly applied to the cell chamber. *In vivo* experiments with macroencapsulated mouse and human islets confirmed insulin delivery efficacy. We also explored the versatility of pressure-based insulin delivery for glycemic control in diabetic mice by varying the duration and intervals of pressure-driven insulin delivery. This approach successfully mimicked the timescale and therapeutic effect of subcutaneous insulin injections or infusions, demonstrating potential for clinical applications.

In this study, our primary focus was on insulin transport, which led us to use a simplified setup that did not include oxygenation enhancements. In our *in vivo* experiments, the islet surface density reached as high as 1600 IEQ/cm^2^. We did not explore key factors critical for the long-term success of macroencapsulated islet engraftment, including maintaining islet health. Looking ahead to future long-term studies, it will be essential to incorporate advanced oxygenation techniques into our design. Potential techniques include the direct injection of oxygen into the device, similar to the βAir device^13,14,31,32^, pump-based extravascular convection-enhanced methods^33^, prevascularization of graft sites^35^, and *in situ* oxygen generation methods^36–41^.

In our theoretical analysis, we explored the application of pressure at levels comparable to normal human diastolic blood pressure and determined that a brief duration of approximately 30 seconds is sufficient. We further assessed the efficacy and safety of our approach by demonstrating that the viability of encapsulated primary human islets over a 5-day period with daily *in vitro* dosing remained similar to that observed through diffusion alone. While this pressure level matches physiological norms and involves only a very short application time—factors that contribute to its safety—we also considered an alternative protocol to further enhance safety margins. Should there be any safety concerns with the application of 11 kPa, our theoretical analysis reveals that reducing the pressure tenfold to 1.1 kPa and extending the application time tenfold to 5 minutes is a viable alternative. This longer duration still aligns well with the initial phase of glucose-stimulated insulin secretion (GSIS). Significantly, this reduced pressure of 1.1 kPa is well below the 20 mmHg (2.7 kPa) typically utilized in the low-pressure mode of machine perfusion for pancreas, which is designed to preserve organ functionality^55^. Considering that machine perfusion generally operates at higher pressures, our alternative setting of 1.1 kPa for 5 minutes should be considered very safe and suitable for future research if needed.

## METHODS

### Relationship between chamber thickness and time required to transport 90% of secreted insulin out of the cell chamber

Solving the Fick’s diffusion equation, the time constant for a one-dimensional diffusion process across a membrane is

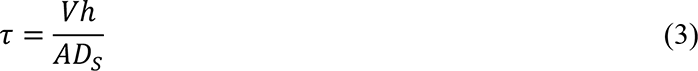

where *V* is the volume of the cell chamber, *A* is the open membrane area, ℎ is the membrane thickness, and *D_s_* is the insulin diffusion coefficient. As *V*/*A* is the thickness of the cell chamber, the diffusion time constant is proportional to this thickness. To transport 90% of secreted insulin out of the cell chamber takes approximately 2.3 time constant.

### Derivation of multiplicative effect of applied pressure relative to diffusion for insulin transport

We start by considering the combined effect of a concentration and a pressure gradient on solute transport across a porous membrane. The combined insulin flux is dependent on flux due to solute diffusional flow, and volumetric solvent flow. This is given by

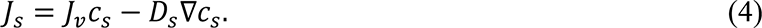

The solute diffusional flow −*D_s_*∇*c_s_* is driven by the insulin concentration gradient, *D_s_* is the solute specific diffusion coefficient, and ∇*c_s_* is the concentration gradient. The insulin flux, *J_v_c_s_*, due to solvent flow is pressure-dependent, with *J*_v_ given by

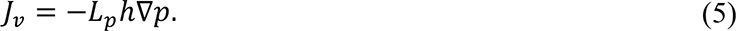

Here, *L_p_* is the filtration coefficient and ℎ is the membrane thickness. We can substitute Eq. 5 into Eq. 4 and yield

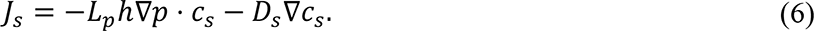

Considering a one-dimensional problem for equation Eq. 6, normalized to membrane thickness, we obtain

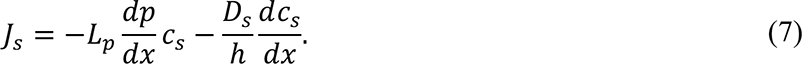

Under steady state conditions, the fluxes due to both pressure and concentration gradients are independent of *x*. This gives a homogeneous second order differential equation:

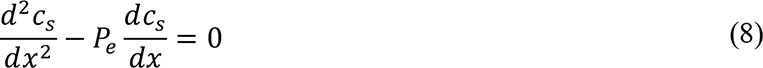

Solving the differential equation in Eq. 8 gives

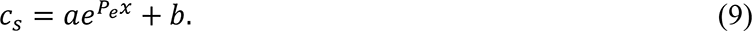

Applying the boundary conditions, we can solve for *a* and *b*:

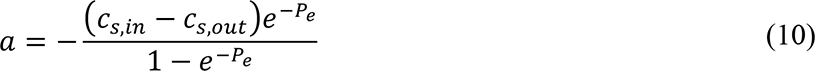

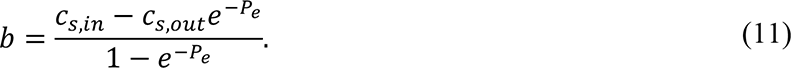

Substituting our solution for *c_s_* to Eq. 7, with 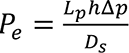 gives

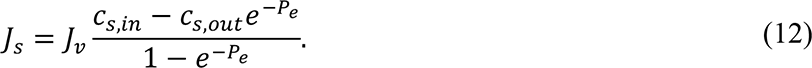

We then take the ratio of combined pressure and diffusion transport to diffusion only transport to yield Eq. 1 in the main text.

### Expressing axial Peclet number in terms of applied pressure and membrane properties

The axial Peclet number is given by

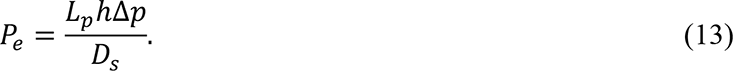

We can apply the Hagen-Poiseuille Law to obtain

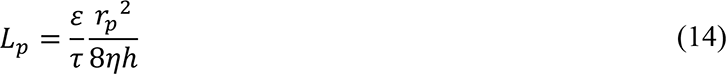

where *ε* is the porosity, *τ* is the tortuosity, and *r_p_* is the pore radius of the membrane, ℎ is the membrane thickness, and *η* is the viscosity of water. For insulin transport where membrane pore radius (*rp*) is much greater than molecular radius of insulin (*rs*), we can apply the Stokes-Einstein equation to obtain

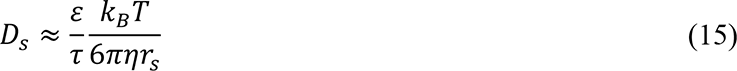

where *k_B_* is the Boltzmann constant and *T* is the absolute temperature. Substituting Eq. 14, 15 into Eq. 13, and simplifying yield Eq. 2 in the main text.

### Design of the PET membrane cell culture insert

Cell culture inserts for 12-well plates were designed (FreeCAD) and 3D printed (Fictiv PLA) (Supplementary Fig. 1). Cell culture inserts included a 9-mm diameter circular opening for membrane attachment, as well as 2-mm diameter tapered circular opening for tubing. PET membranes (Sterlitech PET0413100, PET0213100, PET0813100, Falcon 353180) were attached to the cell culture insert with polyester adhesive (Hotmelt Infinity IM-Supertac-500-12). Wells in a 12-well cell culture plate were filled with 3 mL of media or water.

### Insulin transfer measurements over PET membrane

PET membrane cell culture inserts were placed in a 12-well cell culture plate and loaded with insulin (diluted in Milli-Q water). For diffusion measurements, samples were collected from the fluid in the cell culture plate at time points from 0 to 4 hours. For pressure-based measurements, tubing was connected to a piezoelectric pump, which was used to generate pressure difference across a membrane. Total pump fluid for these experiments was 1 mL. Samples were similarly collected from the cell culture plate fluid at time points from 0 to 30 seconds. The flux was calculated using the 30-second time point. Insulin content was measured by ELISA (ALPCO STELLUX 80-INSMR-CH01).

### R7T1 pseudoislets culture and formation

Immortalized mouse R7T1 β-cells were cultured in DMEM high glucose containing 4 mM L-glutamine and 1 mM sodium pyruvate, with 10% FBS, 1% penicillin/streptomycin, and 1 mg/mL doxycycline (Sigma-Aldrich, St. Louis, MO, USA). R7T1 cells (passage number 10–34) were growth arrested for 5 days with doxycycline-free R7T1 medium supplemented with tetracycline free fetal bovine serum (Gibco A4736301). R7T1 cells were authenticated by immunofluorescence staining (protocol described below) with the following antibodies: insulin (1:500; Cell signaling), Pdx-1 (1:200; Cell signaling), and ZnT8 (1:50; Santa Cruz Biotechnology). To form pseudoislets, growth arrested R7T1 cells were plated at 300,000 cells per well in a 12-well non-adherent plate and after 16–24 hours, pseudoislets were picked and counted for experiments.

R7T1 cells were characterized by comparing growth arrested and proliferating insulin secretion. Cells were growth arrested for 5 days by removing doxycycline from culture media. Proliferating cells were maintained in R7T1 media supplemented with doxycycline for the five-day period. In a 12-well cell culture plate, 300,000 cells per well were plated and cultured in DMEM low glucose (with or without doxycycline) for 24 hours. Media was then replaced with high glucose media and supernatant samples were collected after 30 minutes. Insulin content in supernatant was evaluated by rodent insulin ELISA (ALPCO STELLUX 80-INSMR-CH01).

### *In vitro* encapsulated cell experiments

To measure insulin levels *in vitro*, approximately 800 R7T1 pseudoislets (∼685 IEQ) were loaded onto pre-wetted PET-cell culture inserts. For KCl stimulation experiments, potassium chloride was dosed inside the cell culture insert to achieve a final concentration of 40 mM. After 15 minutes, media samples were collected from inside encapsulation to measure total secreted insulin. Diffusion samples were collected from the media in the cell culture plate. For insulin transport by a brief pressure, a piezoelectric pump and controller were connected to the cell culture insert, and were turned on for 30 seconds, using a total volume of 1-mL, with pressure of 11 kPa. After the pump was turned off, a sample was collected from the media in the cell culture plate. The volume of media outside encapsulation was also measured. Insulin concentration was measured by ELISA (ALPCO STELLUX 80-INSMR-CH01), and total transported insulin was calculated. For non-stimulated experiments, samples were collected after 8 hours of islet encapsulation, and volume was measured. The cell culture insert was moved to a well with fresh DMEM, and the same protocol for insulin transport was followed 2 additional times.

### Human islet information and culture

Human islets were provided by the Alberta Diabetes Institute (ADI) Islet Core (Pro00013094) or the Integrated Islet Distribution Program (IIDP), and used as approved by the Stanford University Administrative Panel on Biosafety. Human islets were cultured overnight in islet medium before experiments. Donor information is provided in Supplementary Tables 1 and 2.

### Islet viability experiments

Human islets (200 µL, 667 IEQ/cm^2^) were encapsulated and cultured in 12-well plates for a period of 5 days. The pressure treatment group was treated with 11 kPa of applied pressure for 30 seconds every 24 hours. To assess islet viability, encapsulated human islets in the diffusion or pressure group were collected, trypsinized, and stained with trypan blue (Corning) at various time points. Cell viability was measured on day 1, 3 and 5, and normalized to day 0.

### Mouse islet isolation

Primary mouse islets were isolated from WT C57BL/6J mice (6–12 weeks of age) by pancreatic perifusion as previously described^56^. Briefly, mouse pancreata were perfused with Cizyme RI (VitaCyte), digested at 37°C for 13 minutes, and islets were purified in 5:6 mixture of Histopaque®-1077 and 1119. Isolated islets were collected from the interface, filtered through a 70-µm cell strainer, and cultured overnight in islet medium (5.6 mmol/L DMEM low glucose containing 4 mmol/L L-glutamine and 1 mmol/L sodium pyruvate, with 10% FBS and 1% penicillin/streptomycin) before selection for experiments.

### Histological analysis

Mouse skin tissue surrounding the implant was harvested, fixed in 4% paraformaldehyde overnight and embedded in paraffin. Deparaffinized and rehydrated tissue sections (5 μm) were stained with hematoxylin and eosin (H&E) or Masson’s trichrome. For immunofluorescence staining, processed tissue sections were antigen retrieved with citrate buffer (pH 6.0) in a pressure cooker for 10 min, blocked with 0.5% donkey serum/0.3% triton X-100/PBS at room temp for 1 hr and incubated at 4℃ overnight with the following primary antibodies: F4/80 (1:500; Proteintech), CD31 (1:500; Proteintech), CD45 (1:500; Proteintech). Antibodies were visualized with secondary antibodies (Jackson Immunoresearch) and imaged with fixed settings on the Leica DMIL inverted fluorescence microscope.

### Non-survival dosing experiments using wild type mice or mice with diabetes

For non-survival experiments, islet insulin was accumulated within the device *in vitro* prior to implantation. Mouse islets, human islets, or R7T1 pseudoislets were suspended in 300-μL DMEM in a non-adherent cell culture dish, supplemented with 1 IU/mL penicillin, 1 µg/mL streptomycin, 1 mM sodium pyruvate, 150 nM Forskolin (R7T1 pseudoislets), and 20 nM exendin-4 (mouse and human islets) for 10 hours. Conditioned media was sampled. Insulin content was measured by ELISA (ALPCO STELLUX 80-INSMR-CH01). The islet/pseudoislet suspension media was then diluted to an insulin concentration of 5 μg/mL. The suspension was mixed thoroughly by pipetting and encapsulated in a 3D printed implant with PET (Sterlitech PET0413100) seal. In experiments involving human islets, 400 human islets were used. Implants were designed (FreeCAD) and 3D printed in PLA (Fictiv). Inserts included a 9-mm diameter circular opening for membrane attachment, as well as 2-mm diameter tapered circular opening for tubing (for pressure dosed implants).

C57BL/J6 mice (10–12 weeks of age) or STZ-induced diabetic C57BL/6J mice (8–10 weeks of age) were anesthetized in 2% isoflurane. Starting blood glucose was 100–180 mg/dL for WT mice, and > 400 mg/dL for diabetic mice. After anesthesia, blood glucose rose to an average of 286 mg/dL in WT mice and 553 mg/dL in diabetic mice. Protectant eye gel was applied. Animals were dosed 500 µL of 0.9% sterile saline intraperitoneally and placed on a 37°C heat source. Breathing rate was monitored throughout the procedure. Animal hair, from the scapulae to the hind quarters, was removed using electric clippers and the shaved area was prepared by wiping with iodine, followed by isopropyl alcohol (repeating 3 times). A ∼1.5-cm incision was made and a blunt probe was used to create space between the skin and the fascia. Blood glucose level was measured every 5 minutes using a glucometer until blood glucose was stabilized (3 consecutive readings with < 10% variation). The implants with encapsulated cells, insulin or sham control media were inserted under the skin and tubing was routed to exit the incision caudally. The incision was closed with surgical staples (Fine Science Tools 12022-09). Insulin dosing was performed immediately after implantation. For pressure-treated cases (and sham control), the tubing was connected to a piezoelectric pump, to generate a pressure difference across the PET membrane of 11 kPa for 30 seconds. No pumping was performed for the diffusion cases. At 0, 15, 30, and 60 minutes, blood glucose was measured using a glucometer and blood samples were collected from the tail vein. Plasma was separated from red blood cells by centrifuging at 10000 RPM. Breath rate was measured at each time point and was > 30 per minute for all mice. After 60–90 minutes, the implant was recovered from the animal. A sample of the conditioned media was collected, to measure residual insulin content. Islets were removed from conditioned media by centrifuging at 2000 rpm for 2 minutes. Plasma insulin content, as well as remaining insulin in encapsulated conditioned media were measured by ELISA (ALPCO STELLUX 80-INSMR-CH01). If animals did not survive, the data was not included in presented results.

### STZ diabetes induction and monitoring

Eight-week-old male C57BL/6J mice were fasted for 4–6 hours. Immediately prior to injection, streptozotocin [100 mM] was suspended in sterile sodium citrate buffer (pH 4.5). Mice were injected intraperitoneally with 175 mg/kg streptozotcin^57,58^ for non-survival experiments, or 150 mg/kg for survival experiments. For three days following injection, drinking water was supplemented with 10% sucrose to prevent hypoglycemia. Once animals reached hyperglycemia (a non-fasting blood glucose > 350mg/dL for several measurements), they were used for insulin dosing experiments. All mice were used for experiments within 1 week of streptozotocin injection. When blood glucose reached >450 mg/dL, mice were dosed 0.05–0.1 unit of insulin subcutaneously daily. The last insulin dose was given no sooner than 14 hours before an experiment (survival or non-survival).

### Survival dosing experiments

For survival experiments, freshly isolated mouse islets were cultured in 1000-μL DMEM in a non-adherent cell culture dish, supplemented with 1 IU/mL penicillin, 1 µg/mL streptomycin, and 1 mM sodium pyruvate for ∼10 hours. The next day, approximately 400 islets were manually picked and moved to fresh DMEM supplemented with 1 IU/mL penicillin, 1 µg/mL streptomycin, 1 mM sodium pyruvate, and 20 nM exendin-4, and introduced into a 3D printed encapsulation implant as previously described. STZ-induced diabetic C57BL/6J mice (8–10 weeks of age) were implanted as previously described. After implantation, animals recovered on a 37°C heat source and were provided with food and water. Blood glucose was measured by glucometer once per hour for 6 hours. After 6 hours, the first insulin dose was delivered. Animals were anesthetized briefly (∼30 seconds). For pressure-driven-dosing animals, and sham controls, the tubing was connected to a piezoelectric pump, which was used to generate a pressure difference across the PET membrane of 11 kPa for 30 seconds. At 0, 15, 30, and 60 minutes and then hourly for 6 hours, blood glucose was measured. At 6 hours, a second dose was provided, as described above, and blood glucose was similarly measured until sacrifice. Animals were provided food and water throughout the full experiment.

In the basal dosing experiments, blood glucose levels were first set to a euglycemic range by injecting a single subcutaneous dose (Humalog 0.05 units). Subsequently, pressure-based dosing was administered for 2.5 seconds every 30 minutes. Throughout the experiment, the animals were freely moving until sacrifice.

### Design of piezoelectric pump controller and wearable wireless micropump

Pizeoelectric micropumps (Servoflo mp6-liq/ mp6-hyb) were used for pressure-driven experiments. A pump driver and controller were designed. An electroluminescent lamp circuit (Microchip HV861) was used as a complimentary dual output piezoelectric driver. Pump rate was calibrated by characterizing the frequency, duty cycle and flow rate relationship. For the wearable wireless micropump, a wireless module (Fanstel BC832) was used to control the piezoelectric driver. Duty cycle was used to adjust the pump rate, and pump time was set by user from a smartphone user interface. A single lithium ion (CR1616) battery was used for power. Silicone tubing (McMaster 2124T2) was connected to the pump, and a reservoir was fabricated out of polyolefin (Pointool PO-01450) and connected to the tubing. Pump fluid consisted of either saline or cell culture media. The power consumption and the weight were optimized to allow both wild type and STZ induced diabetic mice to comfortably carry the full system. The miniaturized system was then packaged in a wearable pouch fabricated out of cotton (Fig. 5E).

Pump pressure was characterized by using a differential pressure sensor (NXP MPX2102DP), with full scale pressure of 100 kPa. Pump outlet was connected to pressure sensor inlet using silicone tubing (McMaster 2124T3). The pump was turned on and the output voltage was measured. Pressure was then calculated based off sensor specifications and measured output voltage. For system pressure measurements, an encapsulation implant was 3D printed (Fictiv PLA) with two openings for silicone tubing. A porous PET membrane was sealed to the open side of the encapsulation implant. One tube was connected to the pump outlet and one tube was connected to the pressure sensor inlet. The pump was turned on, and pressure was measured as described above.

### Statistical analysis

All statistical tests were performed using GraphPad Prism software (GraphPad Software Inc.). Statistical significance was assessed using either Student’s t-test or one-way ANOVA. P-values indicating the significance levels between conditions are reported as \**P* < 0.05, \*\**P* < 0.01, \*\*\**P* < 0.001, and \*\*\*\**P* < 0.0001. Data is represented as mean +/− standard error. The experimental results were independently validated for each dataset presented in this study.

## Acknowledgments

We are thankful to the Alberta Diabetes Institute IsletCore at the University of Alberta in Edmonton with the assistance of the Human Organ Procurement and Exchange (HOPE) program, Trillium Gift of Life Network (TGLN), and other Canadian organ procurement organizations for their support in providing human islets.

## Funding

Stanford Bio-X Interdisciplinary Initiatives Seed Grants Program (IIP) (R10-56) (ASYP, JPA)

IIDP NIH Grant # 2UC4DK098085 (JPA, ASYP)

Stanford SystemX (ASYP)

NIH R01DK101530 (JPA)

NIH R01DK119955 (JPA)

Stanford Graduate Fellowship and Stanford Bio-X Bowes Fellowship (EAT)

Pilot and Feasibility grant from the Stanford Diabetes Research Center, NIH P30DK116074 (SL)

NIH T32 DK007217 (RL, SL)

## Author contributions

EAT performed the experiments, interpreted the data, conceptualized the study, designed the experiments, and wrote the manuscript. SL performed the experiments, designed the experiments, and edited the manuscript. HX performed the experiments, designed the experiments, and edited the manuscript. HM designed the experiments, interpreted the data, and edited the manuscript. JS performed the experiments. RAL conceptualized the study, interpreted the data, and edited the manuscript. JPA conceptualized the study, designed the experiments, interpreted the data, and wrote the manuscript. ASYP conceptualized the study, designed the experiments, interpreted the data, and wrote the manuscript.

## Competing Interests

EAT, RAL, JPA, and ASYP are co-inventors of a patent covering this study filed by Stanford University.

## Data and materials availability

All data are available in the main text or the supplementary materials.

## Notes

### Summary of Updates

We revised the title of the paper to, Enhancing therapeutic insulin transport from macroencapsulated islets using sub-minute pressure at physiological levels, to more accurately reflect the findings we are presenting. In the introduction section, we discuss why current approaches and clinical trials have not achieved insulin independence, highlighting that most strategies prioritize enhancing cell viability to maximize islet surface density. This often involves improving oxygen and nutrient transport from outside to inside the encapsulation, while the transport of insulin from inside to outside has received less attention. To illustrate this point, we have incorporated a new figure (Figure 1), showing that despite these methods improving islet viability within subcutaneous macroencapsulation devices, insulin transport still predominantly depends on diffusion. This introduction provides a foundation for examining the limitations of insulin transport in existing subcutaneous methods. Our focus on insulin transport prompted the use of a simplified experimental setup that excluded oxygenation enhancements, a choice we clarify in the discussion section. There, we emphasize that incorporating oxygenation enhancements to maintain islet health will be crucial for the future success of long-term studies. We have clarified the reference to the pressure levels used in our study. In the introduction, we mention that macroencapsulation methods based on vascular perfusion directly utilize the physiological pressure difference between arteries and veins. This led us to investigate whether a brief application of similar pressure levels could significantly enhance insulin transport. Adopting a conservative approach, we utilized the normal human diastolic blood pressure of approximately 10.7 kPa. Our theoretical insights, grounded in an order-of-magnitude analysis, used 10 kPa as a reference, with experimental validations conducted at 11 kPa. This analysis suggests that a sub-minute application of this pressure level is sufficient and emphasizes the brief nature of the application, contrasting with the continuous pressure in vascular perfusion. In the discussion section, we propose an alternative approach: instead of maintaining constant pressure, we match the pressure application duration with the first phase of glucose-stimulated insulin secretion (5 minutes), resulting in a required pressure of about 1.1 kPa. This pressure is considerably lower than the 20 mmHg (2.7 kPa) used in low-pressure machine perfusion to preserve pancreas functionality, thus affirming 1.1 kPa as a very safe level for future research if needed.

