## Supplementary Material for "Enhancing Therapeutic Insulin Transport from Macroencapsulated Islets Using Sub-Minute Pressure at Physiological Levels"

**This PDF file includes:**

Supplementary Figs. 1 to 9

Supplementary Tables 1 to 2

**Other Supplementary Material for this manuscript includes the following:**

Supplementary Movie 1

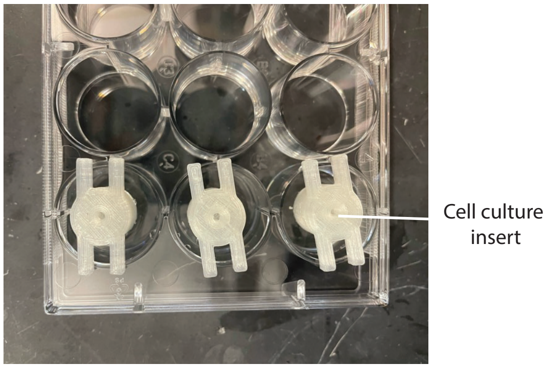


**Fig. 1.** ***In vitro*** **cell culture setup for insulin transport measurement.** The photograph displays the custom cell culture inserts utilized for experimentation in a 12-well plate format, facilitating the measurement of insulin transport across a porous membrane.

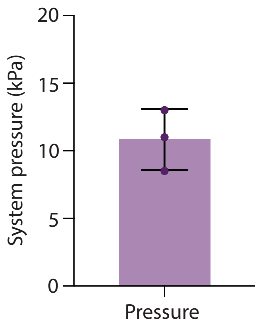


**Fig. 2. Measured pressure at porous membrane driven by the custom-designed piezoelectric micropump system.** Measured pump pressure across three different assembled pump systems.

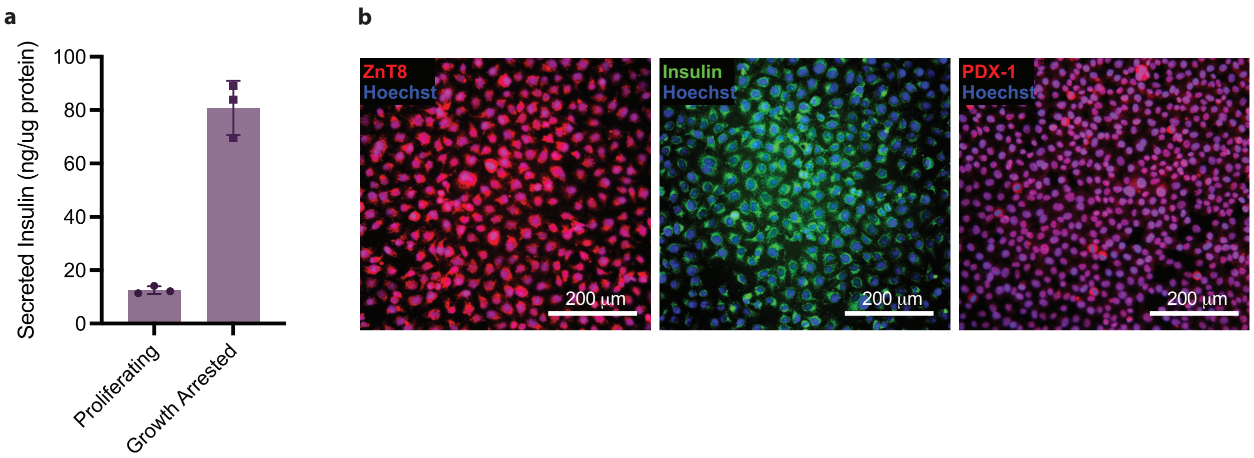


**Fig. 3. Characterization of R7T1 β-cells. a,** Glucose stimulated insulin secretion from growth arrested and proliferating R7T1 cells. Secreted insulin was measured in response to 30 minutes of glucose [25 mM] stimulation after a period of 5 days of growth arrest or non-growth arrested controls, $n = 3$. **b,** Immunohistochemical staining of R7T1 β-cells for identity markers including ZnT8, Insulin, and PDX1.

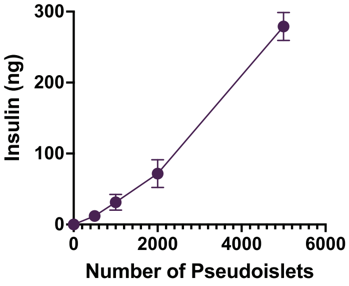


**Fig. 4. Effect of islet number on insulin release.** Insulin released into supernatant by pseudoislet numbers from 0 to 5000 was quantified by rodent insulin ELISA.

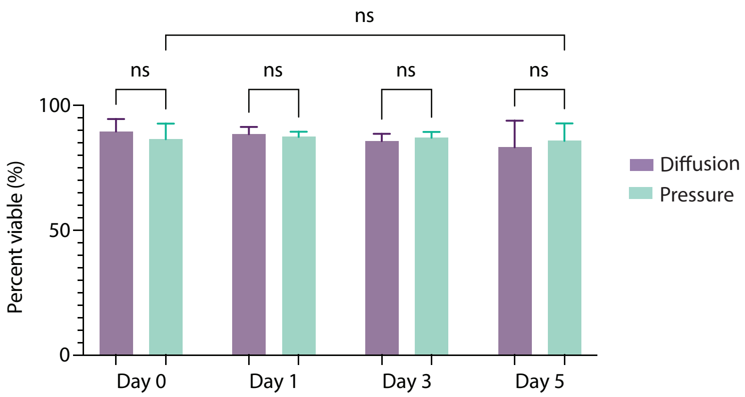


**Fig. 5. Human islet viability in encapsulation with and without daily applied pressure.** Measured viability for a period of 5 days and performed a pressure-driven dose once per day.

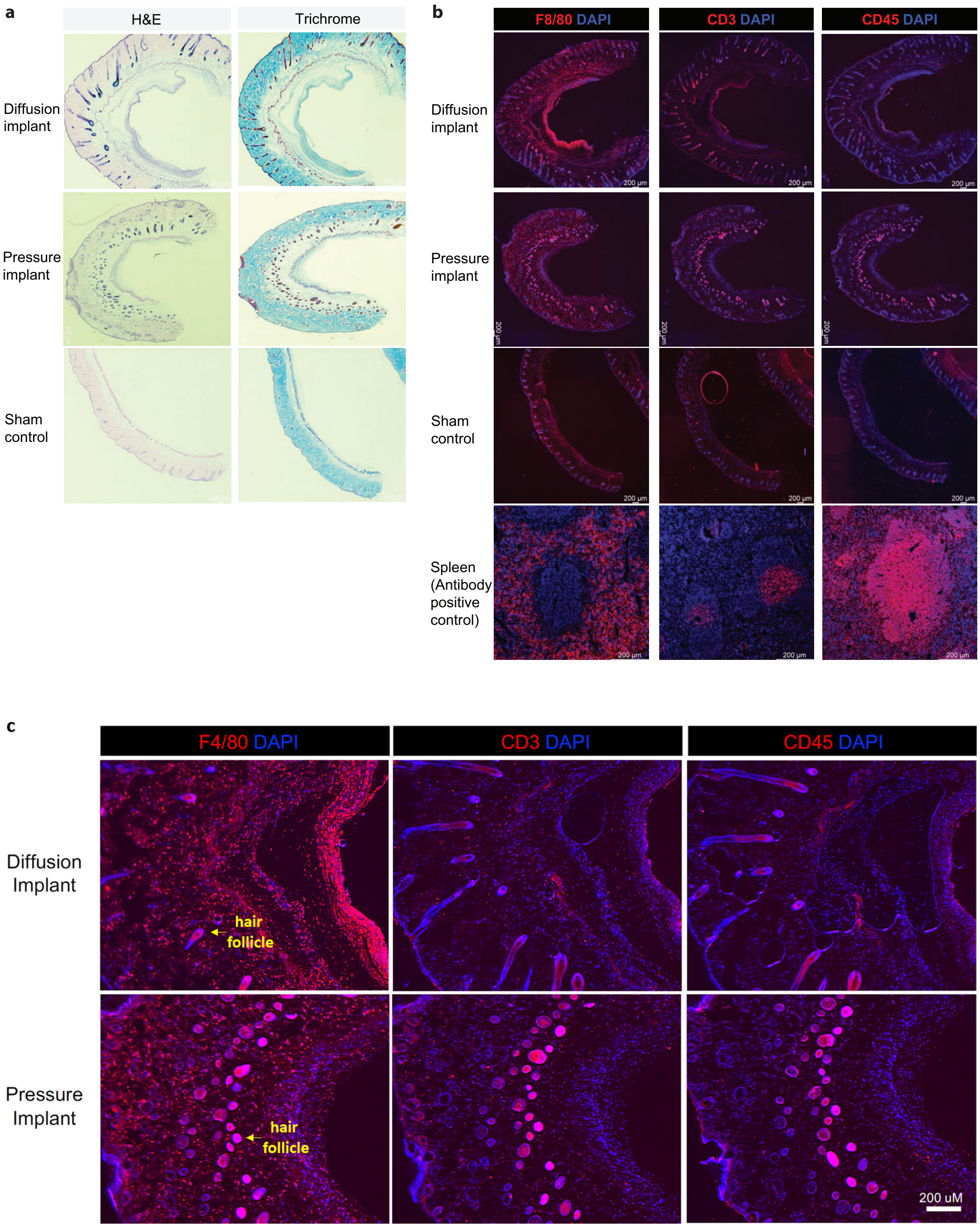


**Fig. 6. Sample histology and immunofluorescence images of tissue sampled from around devices implanted for 14 days.** Insulin-loaded devices were implanted for a period of 14 days.

During this period, devices were used to deliver insulin by diffusion or positive pressure (*n* = 4 each). Subsequently, the devices were extracted and the surrounding tissues (and overlaying skin) were recovered, fixed and sectioned. Healthy skin was also harvested as a control. **a,** Representative images of hematoxylin and eosin (H&E; left) and Trichrome (right) staining. Compared to normal skin, a similar foreign body is observed in association with the device implantation. **b,** Immunofluorescence staining of the same tissues for immunological markers, F4/80 (macrophage), CD3 (T-cell), and CD45 (leukocytes), demonstrates an implantation-associated inflammatory infiltrate. Spleen tissue (bottom row) was included as a positive control for antibody staining. **c,** High magnification images of (**b**) demonstrate substantial macrophage infiltrate with limited T-cell or leukocyte infiltrate. No difference between diffusion- and pressure-treated implants was apparent (*n* = 4 per condition). Background hair root and follicle immunofluorescence is non-specific.

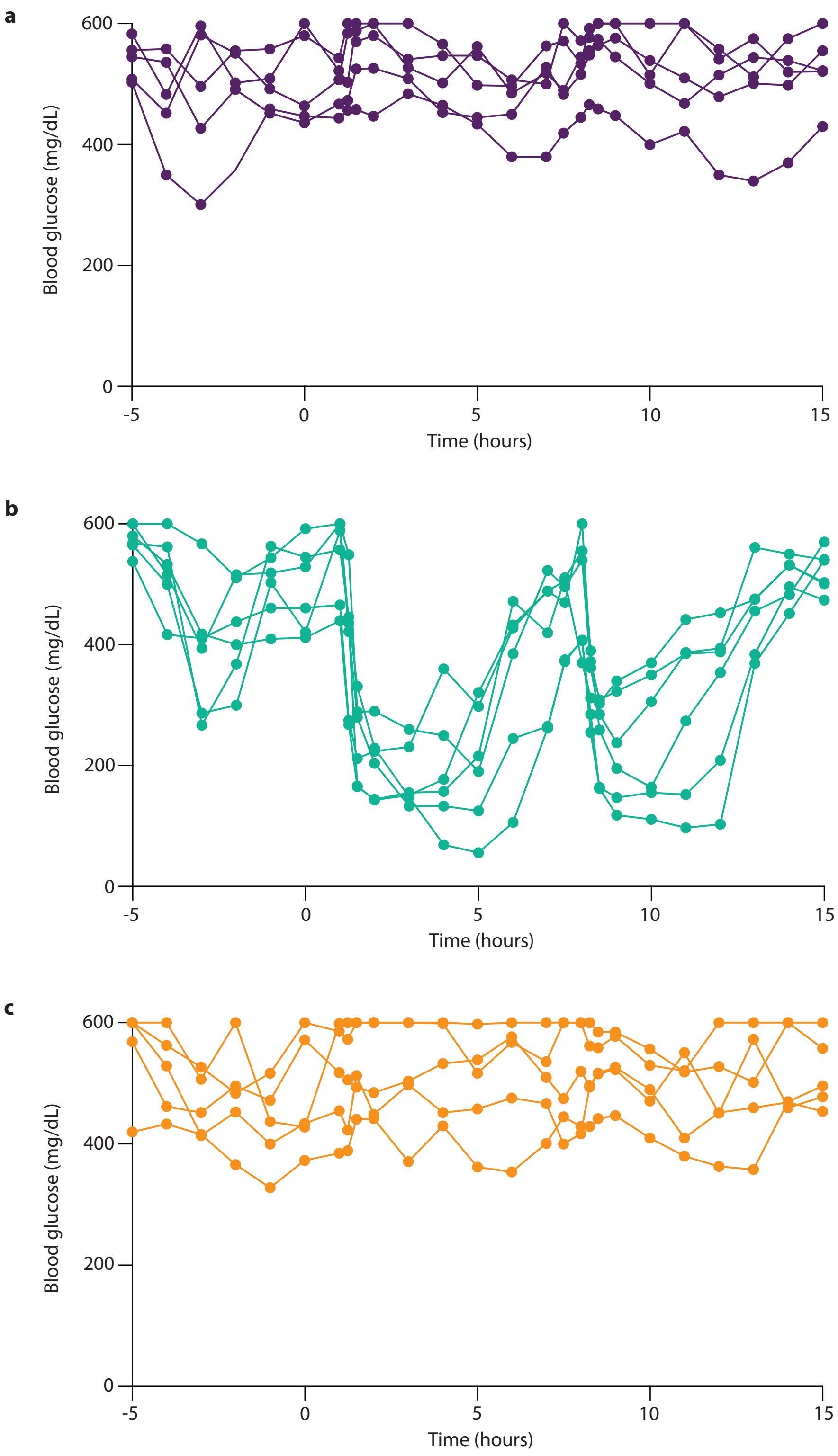


**Fig. 7. Raw blood glucose values from 2-dose insulin boluses experiment in Fig. 6b–d of the main text. a**, Raw blood glucose values for mice treated with insulin dosed via diffusion, $n = 5$. **b**, Raw blood glucose values for mice treated with insulin dosed via applied pressure, $n = 6$. **c**, Raw blood glucose values for sham mice, $n = 5$.

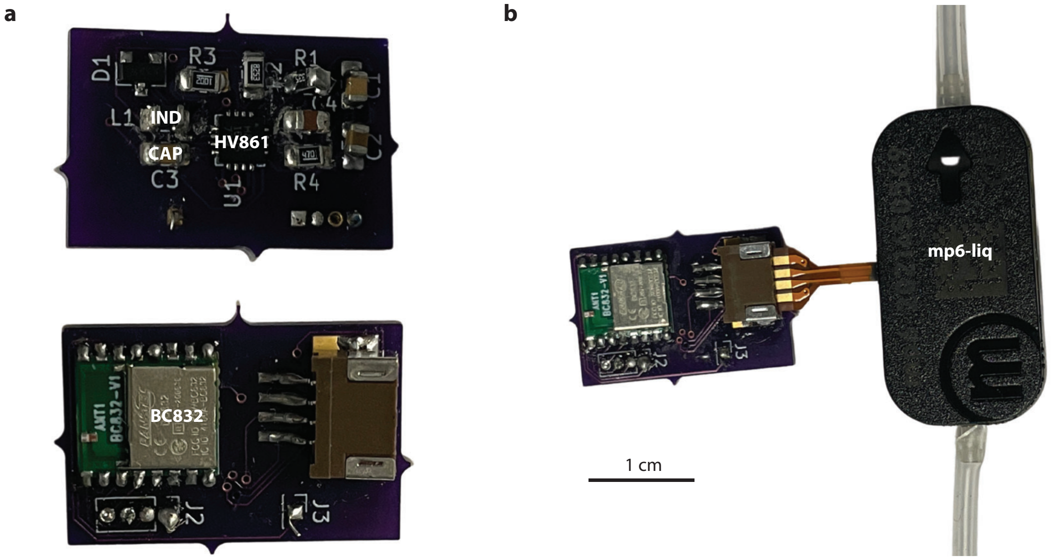


**Fig. 8. Wireless micropump system. a**, Photographs of the front (top) and reverse (bottom) sides of the pump control printed circuit board. The front side includes a high-voltage dual driver (part no. HV861), an inductor (IND), and a few other circuit components for the operation of the driver. The reverse side includes a Bluetooth transceiver module (part no. BC832) and a connector to the piezoelectric micropump. **b**, Photographs of the whole wireless micropump system including the piezoelectric micropump (part no. mp6-liq).

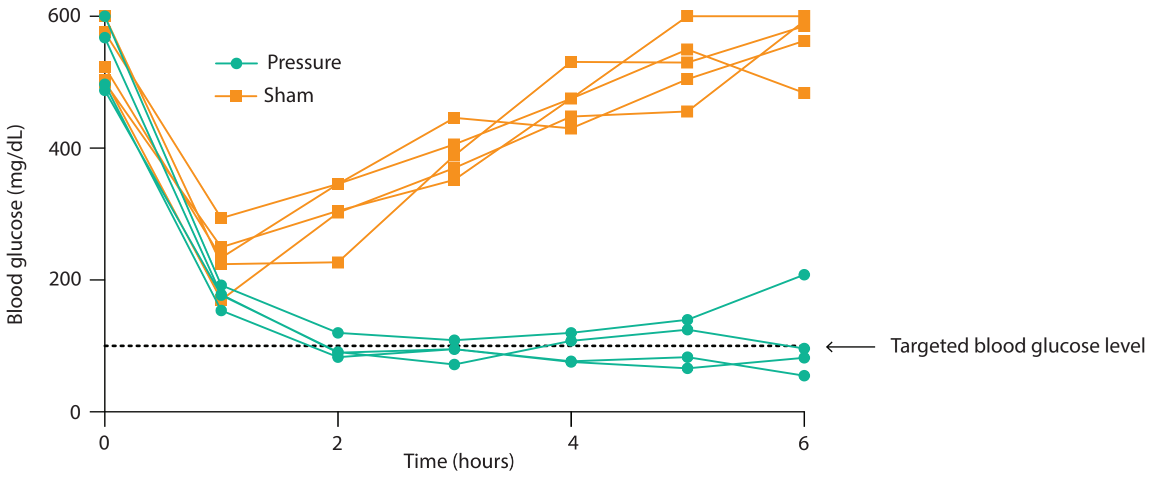


**Fig. 9. Raw blood glucose values from basal dosing experiment in Fig. 6h of main text.** Raw blood glucose values for mice treated with basal insulin dosing. Pressure, $n = 4$; Sham, $n = 5$.

**Table 1: Donor information for the human islets obtained from the Alberta Diabetes Institute (ADI) IsletCore.** HbA1c, glycated hemoglobin; BMI, body mass index; RRID, research resource identified.

| **Donor** | **1** | **2** | **3** |
|---|---|---|---|
| **Gender** | Female | Female | Male |
| **HbA1C (%)** | 5.4 | 5.4 | 5.8 |
| **Age (years)** | 33 | 60 | 55 |
| **BMI (kg/m$^{2)}$** | 31.9 | 25.9 | 23.5 |
| **RRID** | R426 | R421 | R420 |
| **Cold Ischemia Time (h)** | 4 | 18.5 | 3.5 |

**Table 2: Donor information for the human islets obtained from the Integrated Islet Distribution Program (IIDP).** HbA1c, glycated hemoglobin; BMI, body mass index; RRID, research resource identified.

| **Donor** | **1** | **2** | **3** | **4** | **5** |
|---|---|---|---|---|---|
| **Gender** | Male | Male | Male | Male | Male |
| **HbA1C (%)** | | | 4.9 | | 5.4 |
| **Age (years)** | 38 | 52 | 32 | 39 | 16 |
| **BMI (kg/m$^{2)}$** | 28.4 | 37.5 | 26.9 | 32.4 | 29.5 |
| **RRID** | SAMN30181459 | SAMN31242270 | SAMN31815644 | SAMN30686018 | SAMN32641505 |
| **Cold Ischemia Time (h)** | 5.12 | 18.5 | 11.57 | 7.17 | 6.12 |
